# Bodily Maps of Spontaneous Thought

**DOI:** 10.1101/2023.07.06.547921

**Authors:** Hyemin Shin, Byeol Kim Lux, Hong Ji Kim, Choong-Wan Woo

## Abstract

The intricate relationship between the body and the mind has long been recognized, but the specific bodily representations of spontaneous thought remain elusive. Here, we developed and validated predictive models of spontaneous thought based on body maps using the emBODY and Free-Association Semantic tasks. Our valence and self-relevance models demonstrated robust prediction performances across three test datasets, with the valence model accurately decoding the bodily topography of emotions and feelings. Model weight patterns revealed the significance of peripheral limbs and heart area in predicting valence, while the head area played a crucial role in predicting self-relevance. Furthermore, we investigated the neurobiological underpinnings of body map representations using fMRI and ECG data and found evidence for the reflection of body map responses in central and autonomic nervous system activities. Overall, this study provides insights into the bodily representations of spontaneous thought, highlighting the interconnected relationships between the body and the mind.

## Introduction

Following Aristotle’s theory of the “psychosomatic unity,” which posits that the mind and body are inseparable and influence each other^1^, the mind-body relationship has been extensively studied across fields such as psychology, neuroscience, and medicine. Recent research has highlighted the potential role of bodily variables such as interoceptive signals and neuromodulatory systems in evoking and diverting the flow of spontaneous thought^2–5^. Spontaneous thought refers to mental activity that arises in mind with minimal constraints^6–8^, often involving emotional undertones and topics relevant to self-identity^6,9^. It has been suggested that spontaneous thought is an unconstrained memory process in which semantic and episodic memories play an important role^10^, along with bodily sensations^11–13^. In addition, dispositional mind-wandering has been linked to spontaneously arising bodily sensations^14^. Given these findings, it is crucial to consider the relationship between bodily sensations and spontaneous thought when examining the mind-body connection. Here, we aimed to investigate the relationship between spontaneous thought and the body by combining two tasks—the free association semantic task (FAST) for probing spontaneous thought and the emBODY task for measuring conceptual bodily sensations.

The emBODY task has proven to be a valuable tool in revealing the relationship between the body and emotions^15,16^. In the emBODY task, participants are instructed to report bodily sensations in a body-shaped template by coloring the areas they perceived as activated with red and deactivated with blue^15^. These bodily sensation maps appeared to be culturally universal^17^, and the task has shown promise in understanding clinical conditions such as anxiety and depression^18–20^. However, previous studies utilizing the emBODY task have largely focused on discrete emotions, using emotion-related words or exogenous stimuli to trigger bodily sensations. In this study, we employed self-generated stimuli as a more ecologically valid way to induce emotions and provide richer information about the mind-body connection^21,22^. Specifically, we adopted the free association-based thought sampling task, FAST, to elicit self-generated endogenous thought concepts and used the self-generated concepts as stimuli for the emBODY task. The FAST has been previously demonstrated to be a robust tool for probing endogenous spontaneous thought in diverse experimental settings and contexts^23,24^.

In addition, the FAST is uniquely designed to capture detailed information on multiple content dimensions of the self-generated thought concepts, which we leverage in our study to examine bodily sensation maps from a dimensional perspective. The dimensional view of emotions is now widely accepted in the field^25,26^, but previous studies that used the emBODY task have largely focused on investigating the consistency of the body map topography for discrete emotions. By adopting a dimensional approach with predictive modeling techniques, which has shown promise for practical applications^27^, we aim to broaden the scope and enhance the practical utility of body maps.

Furthermore, we collected functional Magnetic Resonance Imaging (fMRI) and electrocardiogram (ECG) data during the FAST task to investigate the neurobiological plausibility of bodily map features. The bodily sensation maps obtained from the emBODY task represent self-reported, and therefore conceptual, bodily activation patterns that potentially include both subjective and biological aspects of our thoughts^16^. Although spontaneous thoughts and emotions are likely to involve bodily sensations, which may be connected to autonomic nervous system activity, such as changes in heart rate^28,29^, there is a lack of evidence that links self-reported thoughts, emotions, and bodily sensations with actual physiological and neural data. Therefore, our novel approach to the emBODY task will contribute to strengthening the neurobiological validity of body map-based studies.

In this study, we aimed to investigate whether conceptual bodily sensation patterns can be used to predict content dimensions of spontaneous thought (**Fig. 1a**). In addition, we sought to evaluate whether the characteristics of newly developed bodily sensation models are consistent with previous studies and whether brain and cardiac activities support the neurobiological plausibility of these models (**Fig. 1a**). To achieve these research goals, we conducted an fMRI experiment with the FAST and a post-scan survey that included the emBODY task on a total of 62 participants (**Fig. 1b**). The experiment consisted of three parts: First, participants generated self-generated concepts in the scanner. Second, they reflected on the concepts while recording ECG activity and undergoing fMRI scanning. Third, in the post-scan survey, they rated each concept on five content dimensions, namely valence, self-relevance, time, vividness, and safety-threat^24,30^, and also they performed the emBODY task for each concept.

**Figure 1.**
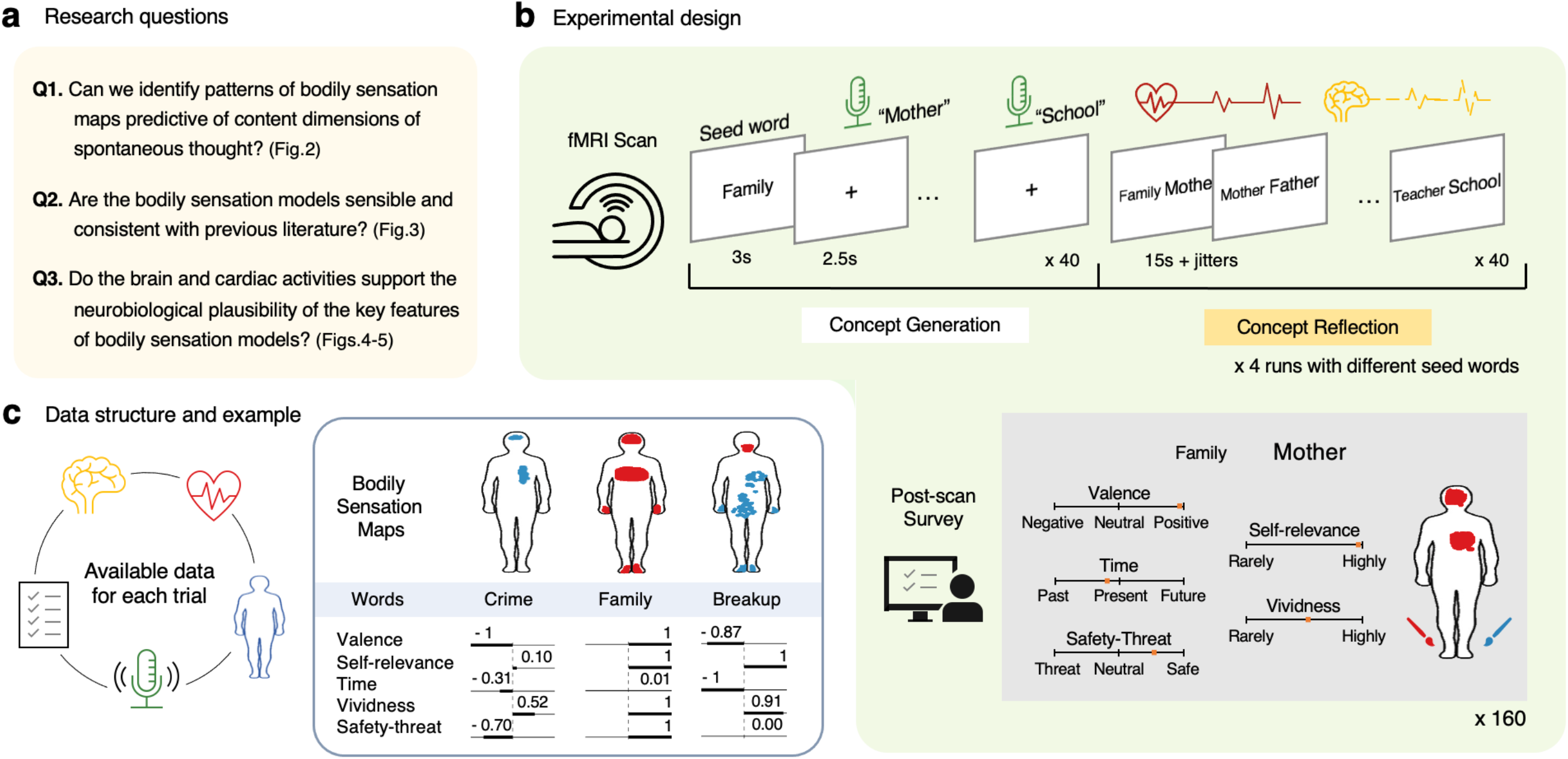
Study overview. **a,** Three main research questions addressed in the study and their corresponding figures. **b,** Experimental design. Participants underwent an fMRI experiment with the Free Association Semantic Task (FAST). During the concept generation phase, participants were presented with a seed word and asked to verbally report a chain of 40 self-generated concepts associated with the preceding response every 2.5 seconds. During the concept reflection phase, participants reflected on each concept of the chain for 15 seconds, with the target concept displayed alongside a preceding concept to provide context for reflection. In the post-scan survey, participants rated each of the 160 self-generated concepts on multiple content dimensions, including valence, self-relevance, time, vividness, and safety-threat. Additionally, they completed the emBODY task for each concept, in which participants reported body areas felt activated in red and areas felt deactivated in blue. **c,** Available data structure and example data. Each trial consisted of a bodily sensation map, heart rate (electrocardiogram) data, a single-trial fMRI activation map (i.e., beta estimate), and rating scores on the 5 content dimensions for each concept. This figure shows three examples of bodily sensation maps and rating scores for the concepts “crime,” “family,” and “breakup.” The bodily sensation maps indicate activated areas in red and deactivated areas in blue. The rating scores ranged from −1 to 1 for valence, time, and safety-threat, and from 0 to 1 for self-relevance and vividness.

We developed body map-based predictive models of each content dimension with machine learning techniques. Our newly developed predictive models demonstrated strong performances in predicting valence and self-relevance across diverse test datasets (**Supplementary Fig. 1**), including external open datasets from the Nummenmaa group^15,16^. Moreover, our results indicated that the body map responses were reflected in both the central and autonomic nervous systems, as evidenced by our analysis of ECG and fMRI data. In particular, we established a relationship between the body map’s heart cluster scores and the heart rate changes measured with ECG. We also observed significant correlations between the heart cluster scores and multiple brain regions within the default mode network (DMN) and hippocampus. Finally, we provided evidence that the body maps conveyed the body-part information that was consistent with the somatotopy of the primary somatosensory cortex (S1).

Our findings provide compelling evidence in support of the validity of the body map-based predictive models, demonstrating their generalizability and neurobiological plausibility. By identifying bodily representations of spontaneous thought, our study findings support the idea of a close connection between the body and the mind, providing a contemporary scientific corroboration for the ancient concept of psychosomatic unity.

## Results

### Data and bodily sensation maps overview

Each self-generated concept was accompanied by a bodily sensation map, heart rate, single-trial fMRI activation map, and rating scores for the five content dimensions (**Fig. 1c**). The reported bodily sensation maps showed activated areas in red and deactivated areas in blue. For example, when a participant reflected on the self-generated concept of “crime,” they reported deactivation in the head and heart areas. Also, the concept was rated as having a negative valence (−1 on a scale of −1 to 1), being self-irrelevant (0.097 on a scale of 0 to 1), related to near past (−0.311 on a scale of −1 to 1), moderately vivid (0.52 on a scale of 0 to 1), and considerably threatening (−0.699 on a scale of −1 to 1).

Across all responses, the most frequently reported body areas included the head, heart, face, and peripheral limbs (i.e., hands and feet), as shown in **Supplementary Fig. 2**. These body areas displayed significant activation and deactivation patterns across participants when averaged and thresholded the maps at *q* < 0.001, false discovery rate (FDR). This suggests that the bodily sensation map data have a low dimensional structure characterized by a limited set of bases. In addition, it highlights the significance of specific body areas, including the head, heart, face, hands, and feet, in representing self-generated concepts.

### Bodily sensation map-based predictive models of spontaneous thought

We developed predictive models based on bodily sensation to answer our first research question (“Can we identify patterns of bodily sensation maps predictive of content dimensions of spontaneous thought?” as shown in **Fig. 1a**), which aimed to identify consistent patterns of conceptual bodily sensations that predict content dimensions of spontaneous thought. We developed the body map-based predictive models using principal component regression with leave-one-subject-out cross-validation (LOSO-CV). As a pre-analysis step, we divided the bodily sensation map data into quartiles based on the trial-by-trial content dimension ratings for each participant and each content dimension. Then, we conducted principal component regression analyses with the preprocessed data. We treated the number of components as a hyperparameter, which was determined based on the cross-validation results with the training dataset (*n* = 29; for details of the cross-validated model performance, see **Supplementary Fig. 3**). We then tested the models on a hold-out test dataset from Study 1 (*n* = 15) and two additional independent test datasets (Study 1 retest, *n* = 19, and Study 2, *n* = 44). The Study 1 retest dataset refers to the data that we collected from a part of the participants of Study 1. For the retest, we used different seed words after an average of 7 weeks. To obtain unbiased prediction performance, we used a cross-validated model that excluded each of the overlapping participants with the training dataset. In addition, Study 2 used a different task parameter, which was a thought-sampling task with a longer interval (50.7 ± 5.6 [mean ± SD] seconds).

Here, we particularly focused on the valence and self-relevance models (**Fig. 2**), as a principal component analysis (PCA) of the five content dimension ratings revealed that the vividness ratings were highly correlated with the self-relevance ratings, and the safety-threat ratings were with the valence ratings (**Supplementary Fig. 4**). Similarly, a PCA on the model weights also showed a similar pattern (**Supplementary Fig. 4**). On the other hand, the predictive model of the time dimension showed poor prediction performance. Hence, we regarded the valence and self-relevance models as our primary focus to examine the connection between the body and spontaneous thought.

**Figure 2.**
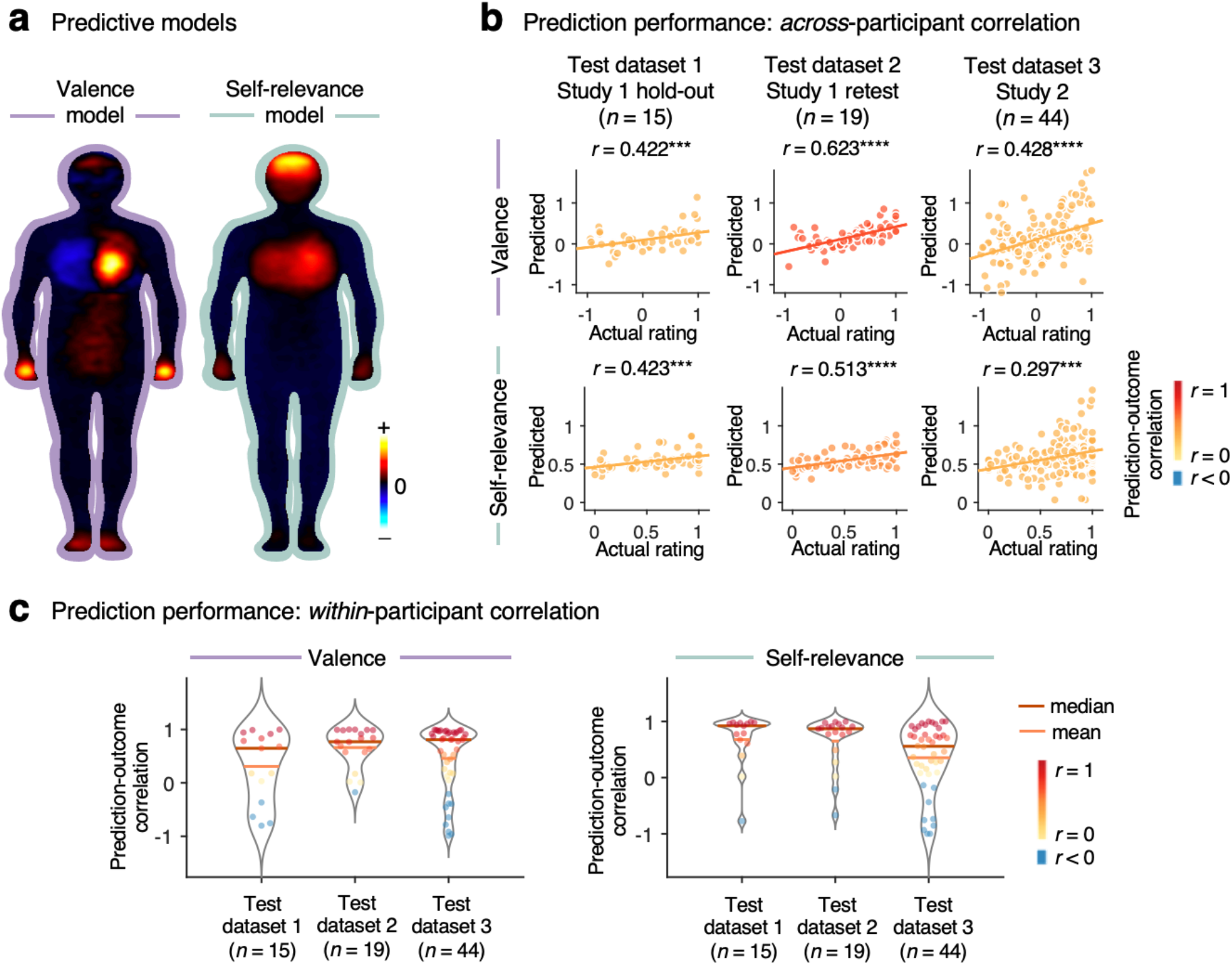
Body map-based predictive models of valence and self-relevance. **a,** Body map-based predictive models of valence and self-relevance. The warm colors indicate positive predictive weights, meaning they positively predict valence or self-relevance scores, while the cool colors indicate negative predictive weights, meaning they negatively predict valence or self-relevance scores. **b,** Prediction performance based on across-participant correlation. The valence and self-relevance models exhibited significantly robust prediction performance across three test datasets. The rows indicate the prediction performance of the valence (top) and self-relevance (bottom) models, while the columns represent different test datasets. From left to right, the plots show the results from 1) the hold-out test dataset of Study 1 (*n* = 15), 2) test dataset 2 of Study 1 retest data (*n* = 19; we conducted cross-validation for testing 11 overlapping participants), and 3) test dataset 3 of Study 2 (*n* = 44). The plots show the correlation between the actual and predicted quartile average scores. Each dot represents the average score of each participant’s quartiles. The line represents the least-square regression line. The colors of dots and lines indicate the level of prediction-outcome correlation. \*\*\**P* < 0.001, \*\*\*\**P* < 0.0001, two-tailed, one-sample *t*-test. **C,** Prediction performance based on within-participant correlation. The within-participant prediction performance ranged from 0.308 to 0.662 for the valence model and from 0.355 to 0.679 for the self-relevance model. Each dot represents the correlation between the actual and predicted quartile averages for an individual participant. The dot colors indicate the level of prediction-outcome correlation. The red and orange lines indicate the median and mean of the participant-level prediction performance, respectively.

The body map-based predictive models of valence and self-relevance are shown in **Fig. 2a**, in which warm colors indicate positive predictive weights (i.e., positively predictive of valence or self-relevance scores) and cool colors indicate negative predictive weights (i.e., negatively predictive of valence or self-relevance scores). The valence model showed high levels of positive predictive weights in the heart, hand, and foot areas, and negative weights in the chest area. The self-relevance model, on the other hand, had high levels of positive predictive weights in the head and chest areas, with moderate levels of positive weights in the hand area. The vividness and safety-threat models had similar patterns to the self-relevance and valence models, respectively (**Supplementary Fig. 5**). As depicted in **Fig. 2b**, the valence and self-relevance models demonstrated good prediction performance across test datasets, with the valence model showing *r*s = 0.422 to 0.623, *P*s = 2.47×10^−9^ to 0.0010, and the self-relevance model showing *r*s = 0.297 to 0.513, *P*s = = 2.56×10^−6^ to 0.0008 (for other models’ results, see **Supplementary Fig. 5** and **Supplementary Table 1**). **Figure 2c** displays the within-subject prediction performance of the valence and self-relevance models, with the averaged correlations between predicted and actual content dimension scores ranging from 0.308 to 0.662 for the valence model and 0.355 to 0.679 for the self-relevance model. All prediction performances of the valence and self-relevance models were significant, except for the test of the valence model on the first test dataset (**Supplementary Table 2**).

We also examined cross-prediction performance to test the specificity of the models. For this, we applied the valence and self-relevance model to predict the other dimension scores on the Study 2 dataset (the third test dataset). The results revealed that the valence model performed better in predicting the valence dimension score compared to the self-relevance model, with *r* = 0.456, *P* = 9.67×10^−7^ for the valence model and *r* = 0.345, *P* = 0.0023 for the self-relevance model, but the difference was not significant, *t*_43_ = 1.5, *P* = 0.1410, paired *t*-test, two-tailed. Similarly, the self-relevance model performed better in predicting the self-relevance score than the valence model, with *r* = 0. 355, *P* = 0.0001 for the self-relevance model and *r* = 0.265, *P* = 0.0068 for the valence model, but this difference was also not significant, *t*_43_ = 0.8762, *P* = 0.3858 (**Supplementary Fig. 6**). These results indicate that the valence and self-relevance models may not be entirely specific to their respective dimensions, possibly because of the common feature of positive predictive weights in the heart area for both models.

### Testing the valence model on external datasets

To assess the sensibility and consistency of our bodily sensation models in relation to prior literature and to address the second research question (“Are the bodily sensation models sensible and consistent with previous literature?” as shown in **Fig. 1a**), we applied our valence model on the datasets derived from studies conducted by the Nummenmaa group^15,16^, which we refer to as Nummenmaa 2014 and Nummenmaa 2018, respectively. Their studies provide patterns of the bodily sensation maps for different types of emotions and feelings, as well as corresponding emotional valence scores. These data enabled us to test our valence model. We applied our valence model to the body maps from prior work, calculating pattern response values via the dot product of the model and the body maps. We then calculated Pearson’s correlation between the pattern responses and the actual *z*-scored valence scores, which we acquired from Nummenmaa 2018^16^. Our valence model effectively predicted the valence rating scores for numerous emotion and feeling concepts, *r* = 0.624, *P* = 0.0402 for Nummenmaa 2014^15^, *r* = 0.402, *P* = 0.0022 for Nummenmaa 2018^16^, two-tailed, one-sample *t*-test, as shown in **Fig. 3a** and **Supplementary Fig. 7**.

**Figure 3.**
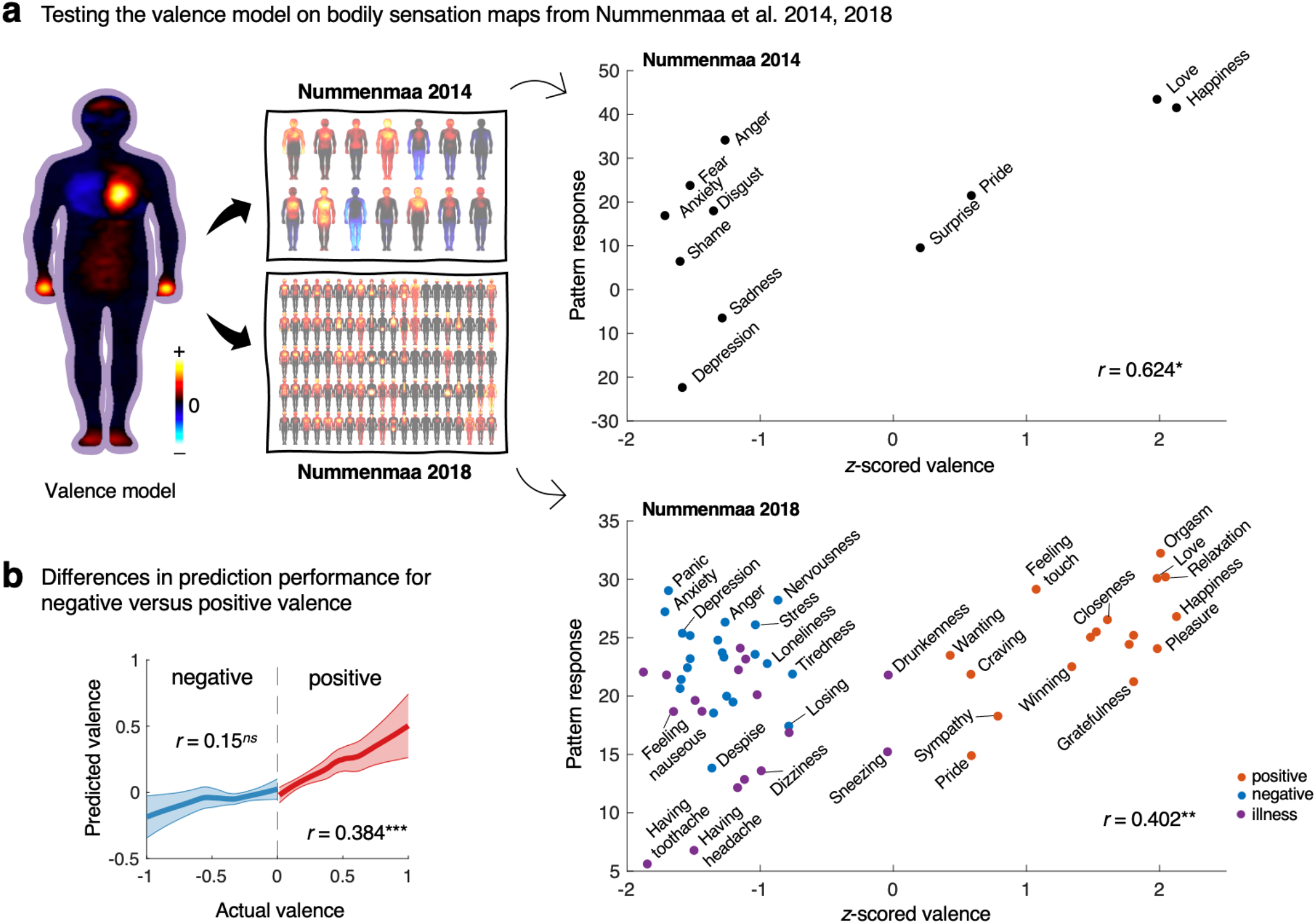
Evaluation of valence model on bodily sensation maps from previous literature. **a,** To evaluate the sensibility and consistency of our valence model with previous research, we tested it on the bodily sensation maps obtained from Nummenmaa et al.^15,16^. The scatter plots in the right panel depict the actual z-scored valence ratings from a previous study^16^ against the pattern responses of the valence model. The pattern response was calculated with the dot product of our model and the body maps from the previous studies^15,16^. Our valence model successfully predicted the valence rating scores for various emotion and feeling concepts. The correlations between the predicted and actual valence ratings were significant for Nummenmaa 2014^15^ (*r* = 0.624, *P* = 0.0402) and Nummenmaa 2018^16^ (*r* = 0.402, *P* = 0.0022). The dot colors in the bottom right panel indicate the clusters of emotions and feelings according to Nummenmaa 2018^16^. \**P* < 0.05, \*\**P* < 0.01, two-tailed, one-sample *t*-test. **b,** The scatter plots in Fig. 3a exhibited a more pronounced linear trend in positive valence compared to negative valence. Motivated by this observation, we further investigated the performance of our valence model for positive and negative valence separately. The plot presents the locally estimated scatter plot smoothing (loess) regression line, depicting the predicted valence plotted against the actual valence scores. Regression analysis was performed separately for data with negative (blue) and positive (red) valence. The prediction results from our test dataset 3 (Study 2) revealed that the model performed better for positively valenced concepts, showing the consistency of the results obtained from the external datasets and our own dataset (*r* = 0.384, *P* = 0.0001 for positively valenced concepts; *r* = 0.150, *P* = 0.1876 for negatively valenced concepts). The line represents the mean of bootstrapped results with 1,000 iterations, and the shading indicates the standard deviation. The correlation values indicate the level of prediction-outcome correlation. *^ns^ P* > 0.05, \*\*\**P* < 0.001, two-tailed, one-sample *t*-test.

However, when we evaluated our prediction based on the positive emotion, negative emotion, and illness clusters provided by Nummenmaa 2018^16^, our valence model performed well only for the positive emotion cluster, *r* = 0.639, *P* = 0.0058, but poorly for the negative emotion and illness clusters, *r* = −0.174, *P* = 0.4391 for negative emotion; *r* = 0.137, *P* = 0.6006 for illness. A similar tendency was also observed in the test results of Nummenmaa 2014^15^. Moreover, further analyses on our third test dataset (Study 2) suggested that the model performed better for positively valenced concepts, *r* = 0.384, *P* = 0.0001; for negatively valenced concepts, *r* = 0.150, *P* = 0.1876, as shown in **Fig. 3b**. These findings indicate that our valence model works better for positive valence, which is consistent across results from external and our own datasets.

### Neurobiological plausibility of the key features of the bodily sensation models

To examine the neurobiological plausibility of our body map-based models, specifically in relation to the third research question (“Do the brain and cardiac activities support the neurobiological plausibility of the key features of the bodily sensation models?” in **Fig. 1a**), we investigated the relationship between the trial-by-trial body maps and cardiac and brain activities measured with an electrocardiogram (ECG) and fMRI. As a pre-analysis step, we performed *k*-means clustering on the trial-by-trial body maps to partition body parts. Then we calculated trial-level bodily sensation scores for the following five body clusters—heart, head, face, hands, and feet, which were among the key features of the valence and self-relevance models. Note that the heart cluster included three sub-clusters.

As the first analysis, we examined the link between cardiac activities while reflecting on self-generated concepts in the scanner and the heart cluster (HC) scores from the post-scan survey (*n* = 50; **Fig. 4a**). We first divided the trials into two groups, high versus low HC trial groups, for each participant based on the median HC score, and then compared heart rate changes from the baseline between two trial groups using paired *t*-tests (**Fig. 4b** left). The results showed that the high HC trials showed increased heart rates compared to the low HC trials across participants at multiple time points throughout the trial (significant at uncorrected *P* < .01 or *P* < .05 as shown in **Fig. 4b**). This indicates that when participants reflected on self-generated concepts associated with higher HC scores, they exhibited an elevated heart rate compared to concepts associated with lower HC scores, suggesting that the self-reported body map could reflect the actual physiological changes beyond mere conceptual bodily sensations.

**Figure 4.**
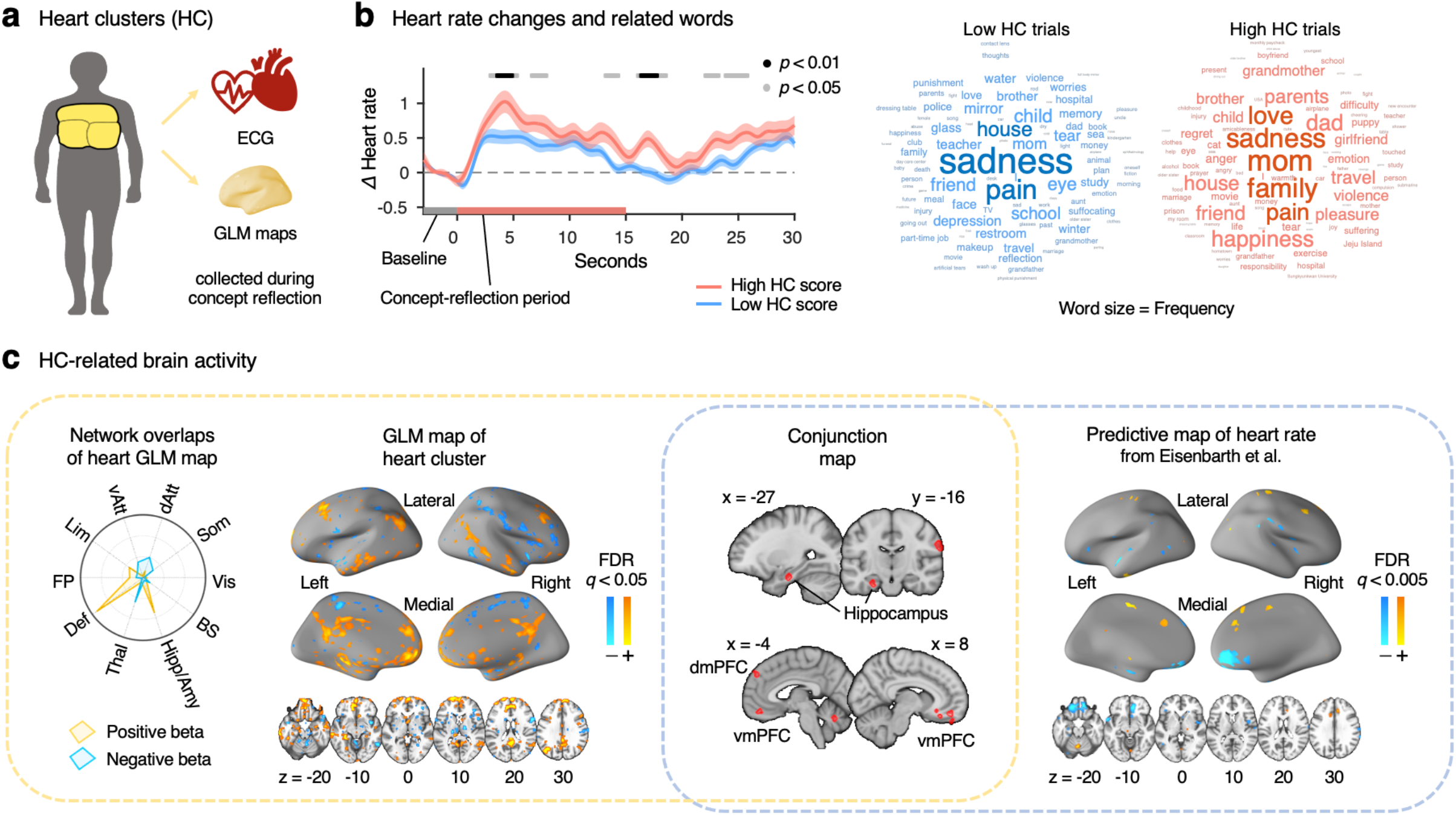
Neurobiological plausibility of the body map’s heart area. **a,** Heart cluster (HC), the body area of interest. The heart cluster included three sub-clusters identified with *k*-means clustering of the bodily sensation maps. **b,** Left: Heart rate changes in beats per minute (BPM) were measured using an electrocardiogram (ECG). The results revealed significantly increased heart rates in trials with high HC scores (red) compared to trials with low HC scores (blue) at multiple time points. The plot represents the average heart rate changes across participants, with the shading indicating the standard error of the mean (Study 1, *n* = 50). Gray and black points on the top of the plot indicate time points at which the high HC score group had significantly higher heart rate changes compared to the low HC score group at uncorrected *P* < .05 and *P* < .01, two-tailed, paired *t*-tests. Right: Word clouds displaying frequently appearing words in the low HC (blue) versus high HC (red) trials. The size of the words represents their relative frequency in each group. **c,** HC-related brain activity. Using a single-trial general linear model on fMRI data obtained during the concept reflection phase (n = 43), we found significant positive correlations between HC scores and brain regions within the default mode network (DMN), hippocampus, and amygdala. (Far left) The radial plot illustrates the overlapping proportions of the Heart Cluster General Linear Model (HC-GLM) map with *a priori* large-scale functional brain networks and subcortical regions^49–51^. Vis, visual; Som, somatomotor; dAtt, dorsal attention; vAtt, ventral attention; Lim, Limbic; FP, frontoparietal; Def, default mode networks; Thal, thalamus; Hipp/Amy, hippocampus/amygdala; BS, brainstem. (Second from the left) The brain map shows the thresholded GLM map for HC at *q* < 0.05 with a false discovery rate (FDR) correction (Study 1, *n* = 43). We then compared the thresholded GLM map with the multivariate pattern-based predictive map of heart rate from Eisenbarth et al.^34^, shown in the rightmost panel. We used the thresholded predictive map available at https://github.com/canlab/Neuroimaging_Pattern_Masks (thresholded at uncorrected *P* < 0.005). (Third from the left) The conjunction map shows the overlapping regions between the FDR thresholded GLM map and the thresholded predictive map, including the hippocampus, ventromedial prefrontal cortex (vmPFC), and dorsomedial prefrontal cortex (dmPFC). These findings provide evidence for the association between the heart cluster of the body maps and neural systems involved in autonomic regulation.

Additionally, we conducted a qualitative analysis of the concepts generated by participants to explore whether there were any differences in the linguistic and semantic aspects between the high and low HC trial groups (the right panel of **Fig. 4b**). The results revealed that the high HC trial group had “mom” (*n* = 47), “family” (*n* = 40), “sadness” (*n* = 40), “love” (*n* = 35), and “pain” (*n* = 32) as the most frequently reported concepts. In contrast, the low HC trial group had “sadness” (*n* = 101), “pain” (*n* = 67), “house” (*n* = 45), “child” (*n* = 40), and “friend” (*n* = 39) (for more information, please see **Supplementary Fig. 8**). It is worth noting that the words “sadness” and “pain” were frequently reported in both trial groups, implying that the conceptual bodily sensations to certain concepts can be influenced by the context in which they were experienced and the personal meaning that the words hold for the individual.

To further examine the neurobiological plausibility of the HC scores, we conducted a general linear model (GLM) analysis with the single-trial fMRI data obtained from the concept reflection phase by regressing the fMRI data on the trial-by-trial HC scores (*n* = 43). Significant positive correlations were found in multiple brain regions within the DMN, hippocampus, and amygdala (FDR *q* < 0.05; **Fig. 4c** left), which are known to play an important role in autonomic regulation^31–33^. In addition, the thresholded map overlapped with a previously developed predictive map of heart rate^34^ (**Fig. 4c** right) in the ventromedial and dorsomedial prefrontal cortices (vm/dmPFC) and hippocampus (**Fig. 4c** third from the left). Including the remaining body parts other than the heart cluster as covariates did not change the results (**Supplementary Fig. 9**).

In addition to the heart area, we found that the hands, feet, and face clusters were represented in the somatotopy of the primary somatosensory cortex (S1; *n* = 37, **Fig. 5**). By contrasting the feet and the face cluster GLM maps within the S1 (**Fig. 5a**), we observed positive beta values in the lateral part of the S1, and the negative beta values in its medial part (**Fig. 5b**). To further investigate the representation of these body clusters in S1, we parcellated the S1 into 10 sub-regions and examined whether the gradients of the contrast values along the sub-regions followed the somatotopy. The mask IDs were assigned numbers starting from the ventral medial part of the S1, which is related to lower-limb somatotopy (e.g., feet), to the dorsal lateral part, which is related to the upper parts of the body (e.g., hands and face) (**Fig. 5c**). Given recent findings suggesting that the representations of certain body parts are distributed across the S1 rather than localized in a specific sub-area^35^, we anticipated observing gradients of the contrasts along the sub-regions.

**Figure 5.**
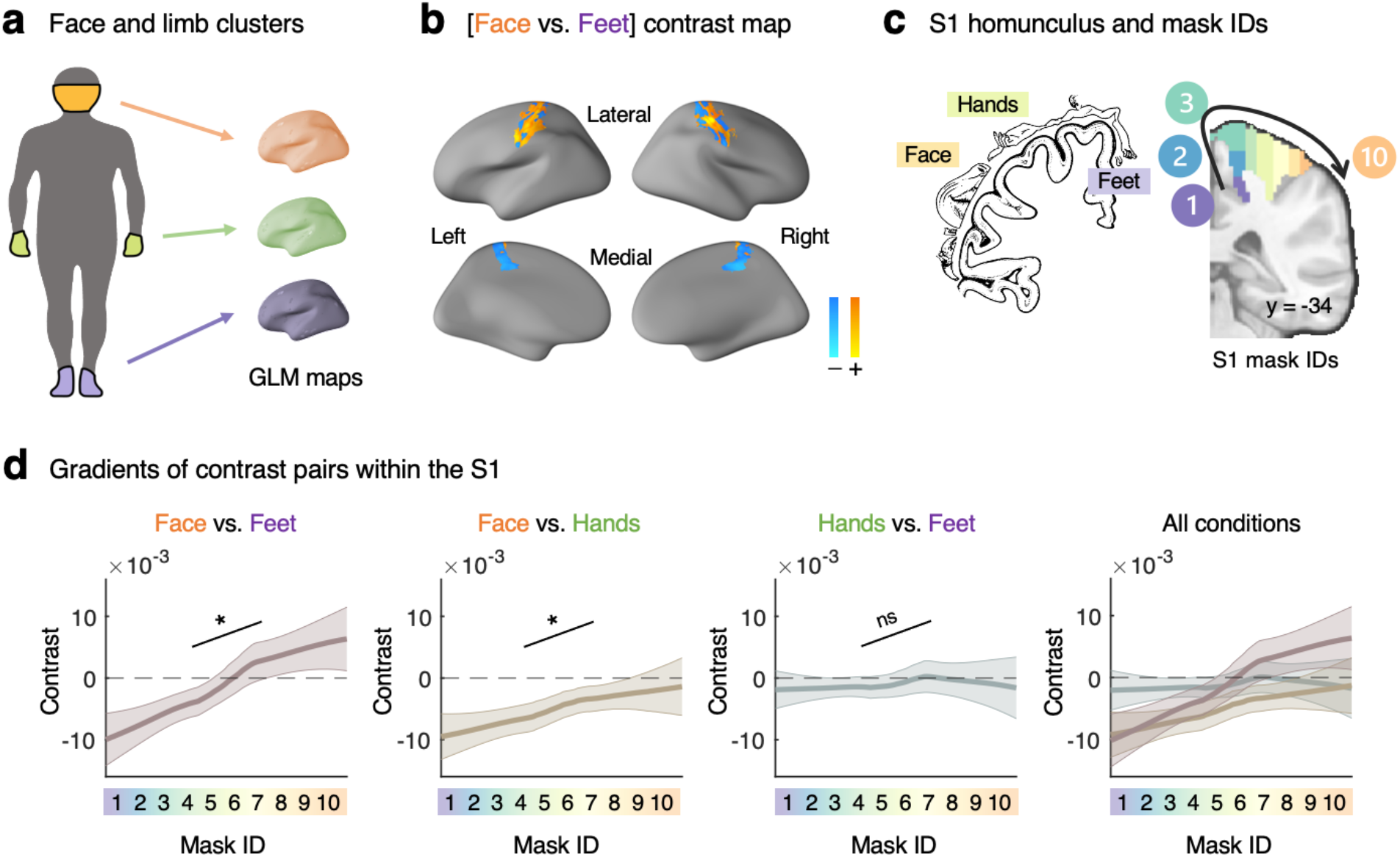
Somatotopic gradients in the primary somatosensory cortex (S1) related to body map clusters. **a,** Face, hands, and feet clusters. **b,** GLM maps for the [face vs. feet] contrast revealed positive betas in the lateral part and negative betas in the medial part of the S1 (unthresholded, Study 1, *n* = 37). **c,** We parcellated the S1 into 10 sub-regions based on the somatotopic gradient, starting from the ventral medial part of the S1, which is related to lower-limb somatotopy (e.g., feet), to the dorsal lateral part, which is related to the upper parts of the body (e.g., hands and face). The schematic S1 homunculus (modified from Fig 17 in the study of Penfield and Rasmussen^52^) depicts the representation of the corresponding body parts, and the assigned mask IDs (1 to 10) within the S1 show their arrangement from medial to lateral regions. **d,** The analysis of the somatotopic gradients for different contrasts revealed that the steepness of the gradients varied depending on the relative distance in the somatotopy between the body parts being contrasted. The [face versus feet] contrast exhibited the steepest slope, followed by the [face versus hands] contrast, and the [hands versus feet] contrast showing a more gradual, non-significant slope. Each plot shows the brain activation gradients for each pair of face, hands, and feet (Study 1, *n* = 37), visualized using locally estimated scatter plot smoothing (loess). The lines represent the mean of bootstrapped results with 1,000 iterations, and the shadings indicate their standard deviations. *^ns^ P* > 0.05, \**P* < 0.05 determined through multi-level GLM analyses with bootstrap tests (10,000 iterations, two-tailed).

The results showed significant gradients across the sub-regions of the S1, ranging from medial to lateral, for both the [face versus feet] and [face versus hands] contrasts (**Fig. 5d**; *z* = 2.51, *P* = 0.012 for [face vs. feet], and *z* = 2.00, *P* = 0.0455 for [face vs. hands]; multi-level GLM, bootstrap tests with 10,000 iterations). Although the gradient for the [hands versus feet] contrast was not significant (z = −0.14, *P* = 0.8857; bootstrap tests with 10,000 iterations), we found that the gradient slopes changed depending on the relative distance between the body parts being contrasted. Specifically, the [face versus feet] contrast, which involves comparing the body parts furthest apart, showed the steepest slope, followed by the [face versus hands] contrast, and the [hands versus feet] contrast displayed the most gradual slope. Overall, our findings provide evidence of a relationship between the body maps and the somatotopic organization of the cortex.

## Discussion

This study examined the bodily representations of spontaneous thought by combining the FAST and emBODY task. We developed body map-based predictive models of valence and self-relevance using machine learning techniques and examined their neurobiological plausibility using fMRI and ECG data. Our main findings are as follows: 1) the patterns of bodily sensation maps can predict the major content dimensions of spontaneous thought, 2) the body map-based predictive models generalize across independent datasets and are consistent with previous literature, and 3) the key features of the models, such as heart, face, and peripheral limbs, display neurobiologically plausible patterns of functional brain correlates and physiological activity. Overall, our results provide evidence that the topography of perceived bodily sensations can account for the multiple dimensions of spontaneous thought and emotions, supporting the interconnected relationship between the body and the mind.

First, the successful development of the body map-based predictive models indicates that the bodily sensation map carries information on emotional valence and self-relevance of spontaneous thoughts. The distinctive and common topographical patterns of the valence and self-relevance models may provide a hint of their neurophysiological underpinnings. For example, the valence model included positive weights in the peripheral limb area, such as hands and feet, which is consistent with a large body of literature suggesting that peripheral sensation and autonomic responses are important sources of emotion^5,21,36–38^. On the other hand, the self-relevance model had positive weights in the head area, implying the dominant involvement of cognitive components such as autobiographical memory and conceptual processing in self-relevant spontaneous thoughts, which is also consistent with previous literature^7,10,39^. In addition, the heart area activation was important for both models, which suggests the centrality of cardiac components (e.g., vagus nerve) in our everyday emotion and cognition^2,40,41^. This common feature of the models could underlie the low specificity observed in the cross-prediction (**Supplementary Fig. 6**).

Second, our models were generalizable across multiple independent datasets, including our own datasets and external open datasets, highlighting their robustness. For example, our multiple test datasets were from different participants and different experimental tasks. The external datasets included the bodily sensation maps of emotions and feeling concepts from two different studies and two different cultures^15,16^. Interestingly, the valence model showed better prediction performance for the positively valenced concepts in both internal and external datasets. When we developed predictive models for positive and negative valence separately by dividing the training data into positive versus negative valence trials, we found that the positive valence model performed well, while it was challenging to develop a well-performing negative valence model (**Supplementary Fig. 10)**. This discrepancy implies that the bodily representations of positively valenced concepts may be relatively simple and linear, whereas negatively valenced concepts may have more intricate representations, as has been noted in previous studies^42,43^.

Lastly, we established the neurobiological plausibility of the body map features using fMRI and ECG data. The GLM analyses showed that the body map scores of the heart area were correlated with the brain activity in the hippocampus, amygdala, and default mode network regions, including vmPFC and dmPFC, which have been suggested to be important for autonomic regulation^31–33^. The scores of the heart cluster in the body map reflected the differences in the heart rate changes measured by ECG. Moreover, the GLM results for the face and peripheral limb clusters in the body map showed the brain activity patterns consistent with the somatotopy in the primary sensory cortex, suggesting that spontaneous thought activated relevant bodily representations in the brain. Notably, the fMRI and ECG data were collected before participants underwent the emBODY task, ensuring that the body map results did not influence the brain and cardiac responses. These results imply that even though the bodily sensation maps are based on self-reports, they can still convey information that has physical relevance.

This study has some limitations that need to be discussed. First, we developed the models using a linear method (here, principal component regression). Although this approach has the advantage of being easily interpretable, it may not capture complex features that could be present in bodily sensation maps. Therefore, future studies could consider using non-linear methods to model body map data. Second, our data appeared to be skewed to the negative valence, possibly due to the self-positivity bias common in healthy populations (**Supplementary Fig. 11**). The self-positivity bias refers to the tendency to report self-relevant aspects (e.g., personality traits, words, etc.) as more positive^44^, which is commonly observed in the healthy population^21,45^. The poor performance of the valence model in predicting negative valence, as well as the poor specificity in predicting valence and self-relevance, could be attributable to this bias. To mitigate this issue, future studies need to include more diverse populations to minimize potential biases in the data. Third, we did not collect data on arousal, a well-known dimension of emotion^46^. Future studies would benefit from investigating the body map representations of arousal as it may be an important factor in bodily sensations. Lastly, we excluded certain participants from specific analyses due to the absence of data (i.e., blank responses) in the emBODY task. This may be ascribed to the unfamiliarity of the task for some participants. To circumvent such issues, it would be helpful to improve task instructions and provide additional practice opportunities.

Despite these limitations, our study broadens the scope of the mind-body connection research by examining bodily sensations associated with spontaneous thought. Bodily topography provides multi-dimensional information that may not be explicitly conveyed verbally. We demonstrated that the bodily sensation maps could capture both the biological and subjective aspects of spontaneous thoughts, thereby providing a more complete picture of our mind. Given that bodily sensation maps, in addition to verbal reports, can provide a more nuanced and comprehensive description of our mind and the body, our findings underscore the potential value of incorporating bodily sensations in emotion research and psychotherapy, as doing so can help uncover hidden aspects of the mind-body connection.

## Author Contributions

B.K.L. and C.-W.W. conceived, designed, and conducted the experiment. H.S. and B.K.L. analyzed the data. H.S. wrote the manuscript. H.S., B.K.L., and C.-W.W. interpreted the results and edited the manuscript. H.J.K. provided the independent test dataset for model testing. C.-W.W. provided supervision.

## Acknowledgments

This work was supported by IBS-R015-D1 (Institute for Basic Science; to C.-W.W.), 2021M3E5D2A01022515 (National Research Foundation of Korea; to C.-W.W.), and Fulbright Foreign Student Program (to B.K.L.).

## Data Availability

The data used to generate figures, including the predictive models, will be shared upon publication through a Figshare repository.

## Code Availability

The codes for generating the main figures will be shared upon publication through a Figshare repository. In-house Matlab codes for fMRI data analyses are available at https://github.com/canlab/CanlabCore and https://github.com/cocoanlab/cocoanCORE.

## Methods

### Participants

We recruited a total of 63 healthy, right-handed participants (mean age = 23.0 ± 2.5 years [mean ± SD], 30 females) for Study 1 from the Suwon area in South Korea. Participants were selected based on predetermined eligibility criteria assessed through an online screening questionnaire, with exclusion criteria applied for individuals with psychiatric, neurological, systemic disorders, and MRI contraindications. A previous publication utilized a subset of the Study 1 dataset^24^, but the analysis results for the bodily sensation map data were not included in that publication. One participant was excluded from the analysis due to an insufficient number of free-association responses for all the analyses. We divided the data of the remaining 62 participants into two subsets: a training dataset for predictive modeling (*n* = 42) and a hold-out test dataset 1 for independent testing (*n* = 20). Additionally, a subgroup of Study 1 participants (*n* = 30, mean age = 22.8 ± 2.3 years, 16 females) underwent a retest session after approximately 7 weeks (51.0 ± 16.8 days). We used this Study 1 retest data as test dataset 2. Furthermore, we recruited 46 participants (mean age = 22.8 ± 2.4 years, 20 females) for Study 2, and their dataset served as test dataset 3. The recruitment of all participants was conducted based on the approval of the institutional review board of Sungkyunkwan University.

From these training and testing datasets, we additionally excluded participants who reported an insufficient number of bodily sensation maps. The exclusion criterion was the blank responses in more than half of the trials. Specifically, we excluded 13 participants from the training dataset, 5 from test dataset 1, 11 from test dataset 2, and 2 from test dataset 3. As a result, the final sample sizes for analysis were as follows: *n* = 29 (mean age = 22.8 ± 2.7 years, 13 females) for the training dataset, 15 (mean age = 22.3 ± 2.1 years, 7 females) for test dataset 1, 19 (mean age = 22.5 ± 2.0 years, 8 females) for test dataset 2, and 44 (mean age = 22.7 ± 2.4 years, 20 females) for test dataset 3 (**Supplementary Fig. 1**). The combined total number of non-overlapping participants included in the various analyses, such as model training, testing, fMRI or ECG analyses, was 100, with *n* = 56 from Study 1 and *n* = 44 from Study 2.

### Experimental design

The Study 1 experiment consisted of three phases: concept generation, concept reflection, and post-scan survey (**Fig. 1b**). In each run of the concept generation phase, participants were presented with a seed word and asked to spontaneously generate a chain of 40 concepts associated with the preceding response every 2.5 seconds. The seed words differed between the first session (“family,” “tear,” “mirror,” and “abuse”) and the retest session (“love,” “fantasy,” “heart,” and “pain”). Prior to the experiment, we conducted a presurvey and selected seed words with an even distribution of the ratings across the valence and self-relevance dimensions. Participants verbally reported a total of 160 concepts across four runs in the scanner.

During the concept reflection phase, participants viewed a pair of consecutive concepts they had generated in sequence while undergoing fMRI scanning. To provide a context for reflection, we displayed a preceding concept alongside the target concept for each trial. The target concept was displayed larger than the preceding one, clearly indicating it as the focus of reflection. We asked participants to ponder the personal meaning of the target concept for 15 seconds, with the stimuli displayed throughout the duration. A fixation cross was presented between trials, with a jittered duration ranging from 3 to 9 seconds.

After the fMRI scanning, the post-scan survey was conducted outside the MR scanner. Participants were asked to rate each self-generated concept on five content dimensions with a continuous scale with a visual analog scale. These dimensions included valence, indicating the positivity or negativity of the concept; self-relevance, measuring the degree of relevance to themselves; time, assessing whether the concept was related to the past, present, or future; vividness, indicating the presence of vivid imagery in the concept; and safety-threat, evaluating the perceived level of safety or threat associated with the concept^24,30^. In addition to the content dimension ratings, participants performed the emBODY task for each concept. During this task, participants used a computer mouse to color body areas where they felt activated in red and areas where they felt deactivated in blue^15^. Activation refers to the feeling of increase (e.g., hot or fast), while deactivation refers to the feeling of decrease (e.g., cold or slow). Participants had the option to use both colors for each concept or to skip coloring if the concept did not elicit any bodily sensations during the reflection process.

Study 2, serving as test dataset 3, comprised an fMRI scan and a post-scan survey. During the fMRI scan, each trial included a free-thinking period lasting an average of 50.7 ± 5.6 seconds, followed by a thought-sampling period of 5 seconds. During the thought-sampling period, participants verbally reported words or phrases that they believed to represent their thoughts from the preceding free-thinking period. In total, the fMRI scan consisted of 42 trials across 7 runs. The post-scan survey included the assessment of the five content dimensions for each concept and the emBODY task, similar to Study 1.

### Body map-based predictive modeling of content dimensions

To develop body map-based predictive models for content dimensions, the bodily sensation map data underwent preprocessing with the following steps: First, we assigned a value of 1 to red-colored body areas and −1 to blue-colored body areas for each trial. Then we convolved and smoothed each body map using a 2-D Gaussian kernel with a σ = 3 pixels. Next, we divided the trial data into quartiles based on the trial-level content dimension ratings. Using the training dataset (Study 1, *n* = 29), we trained predictive models using principal component regression (PCR) with leave-one-subject-out cross-validation (LOSO-CV). We determined the optimal number of principal components based on the cross-validated performance in the training dataset.

To test the generalizability of the developed predictive models, we tested them on the three test datasets. The prediction performances were evaluated using two metrics: 1) across-participant correlation and 2) within-participant correlation between the actual versus predicted rating scores. Predicted rating scores were calculated by summing the intercept and the dot product of the model weights and quartile body map data. Since test dataset 2 (Study 1 retest data) had 11 overlapping participants with the training dataset, we used cross-validated models for testing on those participants.

The body map sizes of Study 2 (537 × 180 pixels) and the external open body map data from Nummenmaa group (522 × 171 pixels) differed from the training dataset (644 × 216 pixels). To match their sizes while preserving aspect ratio, we resized the test data by adding empty pixels and performed up-sampling using bicubic interpolation. For instance, we added two-pixel columns to the left and right sides of the Nummenmaa body maps, resulting in 522 × 175 pixel body maps.

The pattern response scores in **Fig. 3a** were calculated by taking the dot product between the valence model weights and the body maps from Nummenmaa et al. studies. The intercept was not included in the calculation due to the differing scales of the outcome variable across the two studies.

### fMRI data acquisition and preprocessing

The MRI data for Study 1 were collected using a 3T Siemens Prisma scanner at Sungkyunkwan University. High-resolution T1-weighted structural images and functional echo-planar imaging (EPI) data were obtained with a repetition time (TR) = 460 ms, echo time (TE) = 27.2 ms, multiband acceleration factor = 8, field of view = 220 mm, matrix size = 82 × 82, voxel size = 2.7 × 2.7 × 2.7 mm^3^, 56 interleaved slices, and a total of 2608 volumes. Stimulus presentation and behavioral data acquisition were controlled using MATLAB (MathWorks) and Psychtoolbox (http://psychtoolbox.org/).

Data preprocessing was conducted using SPM12 (Welcome Trust Centre for Neuroimaging) and FSL (the Oxford Centre for Functional MRI of the Brain). The structural T1-weighted images were coregistered to the functional images based on the first single-band reference (SBRef) image. The T1 images were then segmented and normalized to the Montreal Neurological Institute (MNI) space. For functional EPI images, the initial 20 volumes were discarded to allow for image intensity stabilization. Outliers were identified and treated as nuisance covariates to remove intermittent gradient and severe motion-related artifacts that were present to some extent in all fMRI data. We detected the outliers using the following criteria: 1) Mahalanobis distances and 2) root mean square of successive differences. Mahalanobis distances were computed for the matrix of concatenated slice-wise mean and standard deviation values over time. Images exceeding 10 mean absolute deviations were identified as outliers using moving averages with a full width at half maximum (FWHM) of 20 images kernel. Outliers based on the root mean square of successive differences across volumes were identified as images exceeding three standard deviations from the global mean.

We then conducted motion correction (i.e., realignment) using the SBRef image as a reference and applied distortion correction using FSL’s topup function. Lastly, the functional images were normalized to the MNI space using the normalization parameters derived from T1 normalization. The interpolation was set to 2 × 2 × 2 mm^3^ voxels, and the images were smoothed with a 5-mm FWHM Gaussian kernel. As the data from two participants had poorer image quality after distortion correction, we used distortion-uncorrected images for these participants.

### fMRI single-trial analysis

For the fMRI analysis in Study 1, we employed a single-trial analysis approach. The estimation of the single-trial response magnitude for each voxel was performed using a general linear model (GLM) design matrix with separate regressors for each trial, following a similar approach to the ‘beta series’ method^47^. To construct the design matrix, the data from the concept reflection phase were concatenated for each participant, with run intercepts added. Regressors were created for the trials of the concept reflection phase using a boxcar convolved with SPM12’s canonical hemodynamic response function (HRF). The design matrix also included various nuisance covariates, such as dummy coding regressors for each run (representing run intercepts), a linear drift across time within each run, 24 head motion parameters for each run (including x, y, z, roll, pitch, and yaw, as well as their mean-centered squared values, derivatives, and squared derivatives), indicators for outlier time points, and the top five principal components of white matter and cerebrospinal fluid signals. We conducted the first-level analysis using the design matrix in SPM12, with a high-pass filter (180 s). To account for potential acquisition artifacts that may occur during trials and significantly affect single-trial estimates, trial-level variance inflation factors (VIFs) were calculated. Trials with VIFs exceeding 3 were excluded from further analyses. On average, the number of excluded trials due to high VIFs was 2.902, with a standard deviation of 3.081.

### Univariate voxel-wise GLM analysis

To investigate the neural correlates of the body map cluster scores, we conducted the general linear model (GLM) analysis using the fMRI data from the Study 1. We had to exclude 19 participants from the dataset of 62 participants due to either insufficient brain coverage in the MRI data (*n* = 1) or an insufficient number of body map responses (i.e., fewer than half of the trials; *n* = 18). Trials with no emBODY task response were also excluded for each participant. In the voxel-wise GLM analysis, the independent variable (x) was the trial-by-trial scores of each body map cluster, while the dependent variable (y) was the single-trial beta maps for the concept reflection phase. To identify body clusters in a data-driven manner, *k*-means clustering was initially performed. We then focused on specific body parts deemed important in further GLM analyses, such as the heart, head, face, hands, and feet. Trial-level body cluster scores were obtained by averaging the body map clusters for each trial. For each GLM analysis of a particular body part, the body cluster scores were z-scored within each participant to normalize them onto the same scale across all participants. Additionally, participants who had zero scores across all trials for the particular body part were excluded from the analysis (*n* = 0 for the heart and face clusters, *n* = 1 for hands, and *n* = 6 for feet). Regarding the GLM analysis of the heart clusters, which consisted of three sub-clusters identified through *k*-means clustering, the beta maps from the three individual clusters were concatenated. Voxel-wise one-sample *t*-tests were then conducted across participants, and the result was corrected for multiple comparisons using a false discovery rate (FDR) threshold of *q* < 0.05.

### Cardiac data analysis

Cardiac data were collected using an electrocardiogram (ECG) during the fMRI experiment in Study 1 (*n* = 62). MR-compatible electrodes, placed at the right and left clavicles and left lower abdomen area (Biopac Systems, Goleta, CA) were used to record the ECG activity. The ECG data were sampled at a rate of 2000Hz. To eliminate MR-related noise, two filters were applied: 1) a band-pass filter ranging from 0.6 to 10 Hz, and 2) a band-stop filter for which we used multiples of 1/TR (1/0.46 Hz) to account for the potential reciprocal influence of MRI sampling frequency. PhysIO Toolbox (https://www.nitrc.org/projects/physio/)^48^ was utilized to identify peaks in the ECG data and calculate the inter-beat interval (IBI) by measuring the distances between the identified peaks. The IBI data were down-sampled to 25 Hz and then low-pass filtered (< 0.5 Hz). Beats per minute (BPM) was calculated by dividing 60 seconds by the IBI (60/IBI). The data from the last trial for each participant were excluded due to incomplete recording. Additionally, the data from 5 participants were excluded due to recording issues, and the data from an additional 7 participants were excluded due to unusual variance (< 0.0001 or > 2) in participant-level BPM data across the time course. This resulted in a final sample size of 50 participants for further analysis. To examine the changes in heart rate during the concept reflection phase, the baseline BPM for each trial was calculated using the mean BPM of the 3 seconds right immediately preceding the stimulus onset. The baseline BPM was then subtracted from the trial-level BPM data to determine the heart rate changes for each trial. To investigate the difference in heart rate changes between trials with high versus low heart cluster (HC) scores, we first divided the trials into two groups of trials based on the median HC score. Subsequently, the BPM data were averaged across the trials within each group (i.e., high versus low HC scores) for each participant.

### Statistical analysis

In **Figs. 2b**, **3a**, and **3b**, we performed two-tailed one-sample *t*-tests to determine if the prediction-outcome correlations (correlations between the predicted and actual rating scores) were significantly different from zero. For **Fig. 2c**, we conducted bootstrap tests with 10,000 iterations (two-tailed) to examine whether the mean of the within-participant prediction-outcome correlations was greater than zero. The sample sizes for the test datasets in **Figs. 2b-c** were *n* = 15, 19, and 44 for test dataset 1, test dataset 2, and test dataset 3, respectively. In **Fig. 3b**, we used test dataset 3 (*n* = 44). In **Fig. 4b**, we conducted two-tailed paired *t*-test at each time point to assess if the heart rate changes between high HC score trials and low HC score trials significantly differed across participants (*n* = 50). In **Fig. 4c**, two-tailed one-sample *t*-tests were applied to the GLM results, and the results were thresholded using false discovery rate (FDR) correction at *q* < 0.05 for visualization purposes (*n* = 43). In **Fig. 5d**, we conducted multi-level GLM analyses with bootstrap tests (10,000 iterations, two-tailed) on the linear slopes to determine if the slopes were significantly greater than zero (*n* = 37).

