## Supplementary information for "Bodily Maps of Spontaneous Thought"

**Running Head:** BODILY MAPS OF SPONTANEOUS THOUGHT

**This PDF file includes:**

1. Supplementary Figs. 1-11

2. Supplementary Tables 1-2

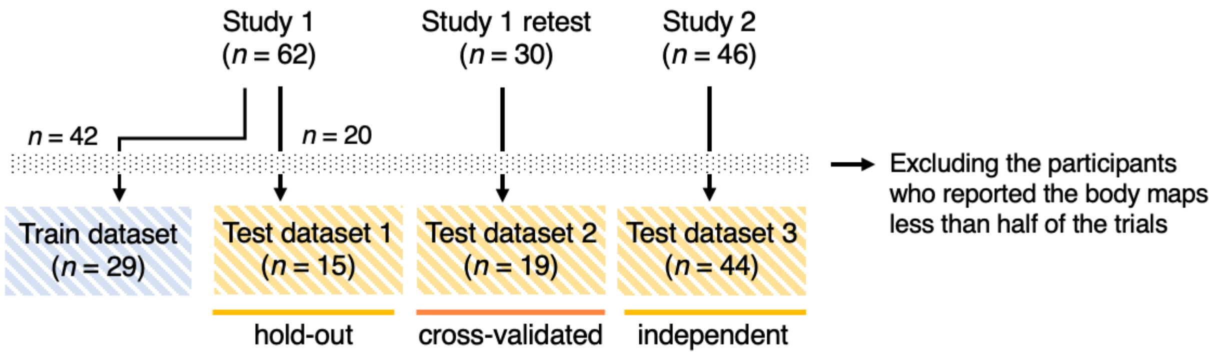


**Supplementary Fig. 1 (related to Fig. 2). Train and test datasets.** We analyzed four different datasets from two different studies; the train dataset and test dataset 1 were from Study 1, and test dataset 2 was from the retest session of Study 1. The test dataset 3 was from Study 2. We excluded the data from participants who reported the body maps in less than half of the trials: For the training dataset, test dataset 1, and test dataset 2, the participants who reported less than 80 out of 160 trials were excluded, and for the test dataset 3, the participants who reported less than 21 out of 42 trials were excluded. The test datasets 1 and 3 were fully independent from the train dataset. For the test dataset 2, we used the cross-validated models, i.e., we tested the models that were trained on the train dataset excluding the overlapping participant's data.

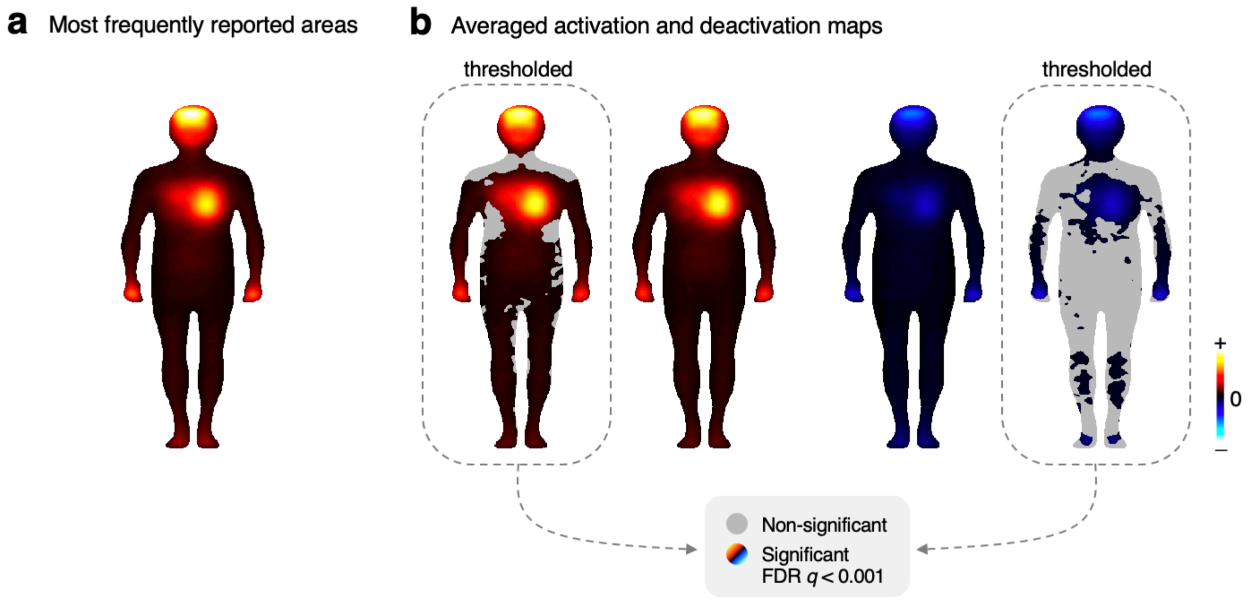


**Supplementary Fig. 2 (related to Fig. 2). Descriptive analysis of the bodily sensation maps.** We conducted descriptive analyses for the body map responses. **a,** To examine the most frequently reported areas, we calculated the averaged absolute values of the trial-by-trial body map responses of Study 1 ($n$ = 62). The head, heart, face, and peripheral limbs (i.e., hands and feet) were the most frequently reported body parts. **b,** The averaged body maps separately for the activation and deactivation across participants (Study 1, $n$ = 62). The thresholded maps show the body areas significant at false discovery rate (FDR) $q$ < 0.001, two-tailed, one-sample $t$-test across participants.

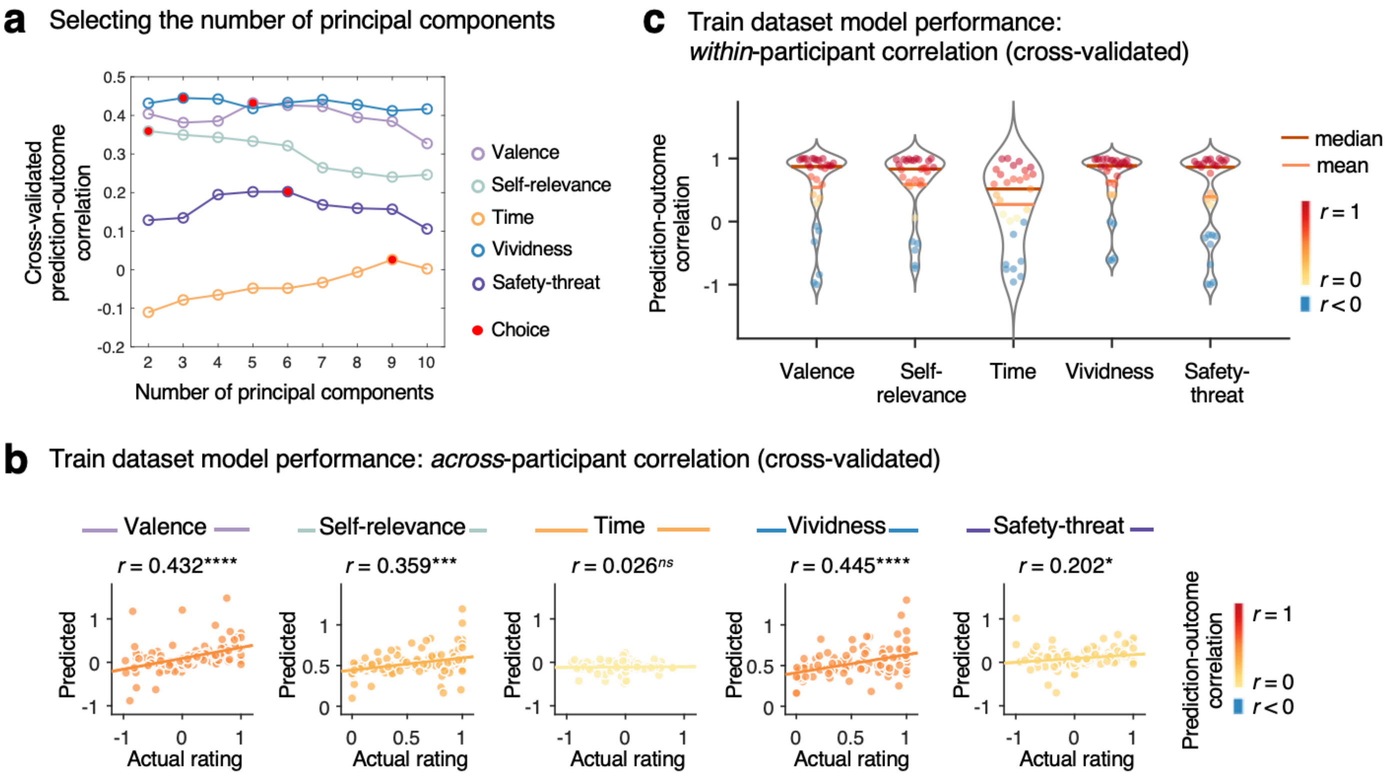


**Supplementary Fig. 3 (related to Fig. 2). Body map-based predictive modeling. a,** The plots show the cross-validated model performance (i.e., across-participant prediction-outcome correlation) for different numbers of principal components (PCs). Lines with different colors indicate the results for different content dimensions. The red dot indicates the selected number of PCs that provided the best cross-validated model performance (5 PCs for valence, 2 PCs for self-relevance, 9 PCs for time, 3 PCs for vividness, and 6 PCs for safety-threat). **b,** The cross-validated across-participant prediction performance in the train dataset for each dimension, for valence, $r = 0.432$, $p = 2.15\times10^{-6}$; for self-relevance, $r = 0.359$, $p = 0.0001$; for time, $r = 0.026$, $p = 0.7856$; for vividness, $r = 0.445$, $p = 9.85\times10^{-7}$; for safety-threat, $r = 0.202$, $p = 0.0349$; two-tailed, one-sample $t$-test, $^{****}p < 0.0001$, $^{***}p < 0.001$, $^{*}p < 0.05$, $^{ns}p > 0.05$. Scatter plots show the correlation between the actual versus predicted quartile average scores. The line represents the least-square regression line. **c,** The cross-validated within-participant prediction performance in the train dataset for each dimension. The mean prediction-outcome correlation was for valence, $r = 0.541$, $p = 2.63\times10^{-6}$; for self-relevance, $r = 0.586$, $p = 7.46\times10^{-9}$; for time, $r = 0.271$, $p = 0.0212$; for vividness, $r = 0.643$, $p = 5.92\times10^{-12}$; for safety-threat, $r = 0.392$, $p = 0.0020$; bootstrap tests with 10,000 iterations. Each dot represents the correlation between the actual versus predicted quartile averages for each participant. The red and orange lines indicate the median and mean of the participant-level prediction performance, respectively.

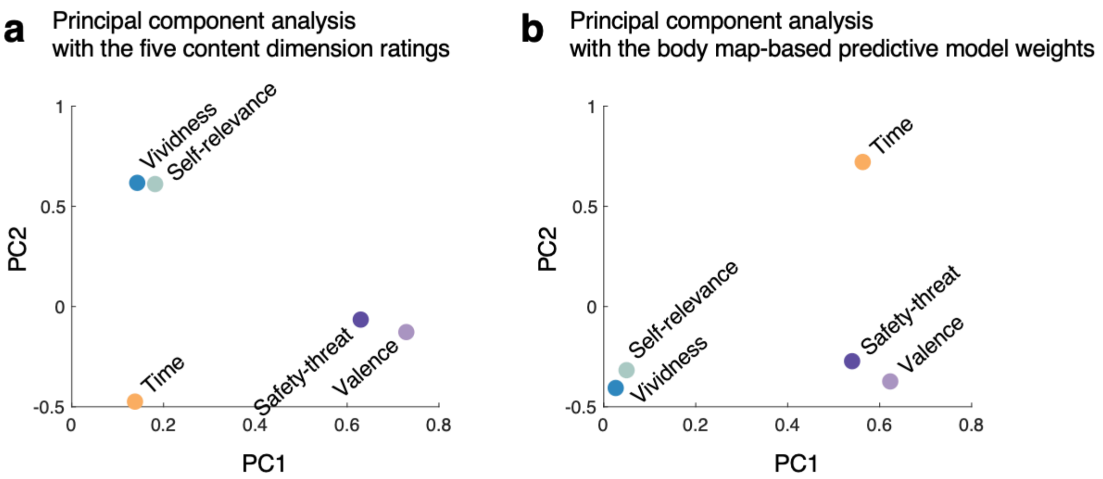


**Supplementary Fig. 4 (related to Fig. 2). Principal component analyses on the ratings and models.** We conducted the principal component analyses (PCA) using the Study 1 data ($n$ = 62) and body map-based predictive model weights. **a,** The plot shows the PCA loadings based on the trial-by-trial five content dimension ratings. When we plotted the PCA loadings on the top two principal component axes, the self-relevance and vividness were located in a similar position, and the valence and safety-threat dimensions were close to each other. The time dimension was located far from the other two clusters. **b,** The plot displays the PCA loadings based on the body map-based predictive model weights. The patterns were similar to the results based on the ratings in **a**.

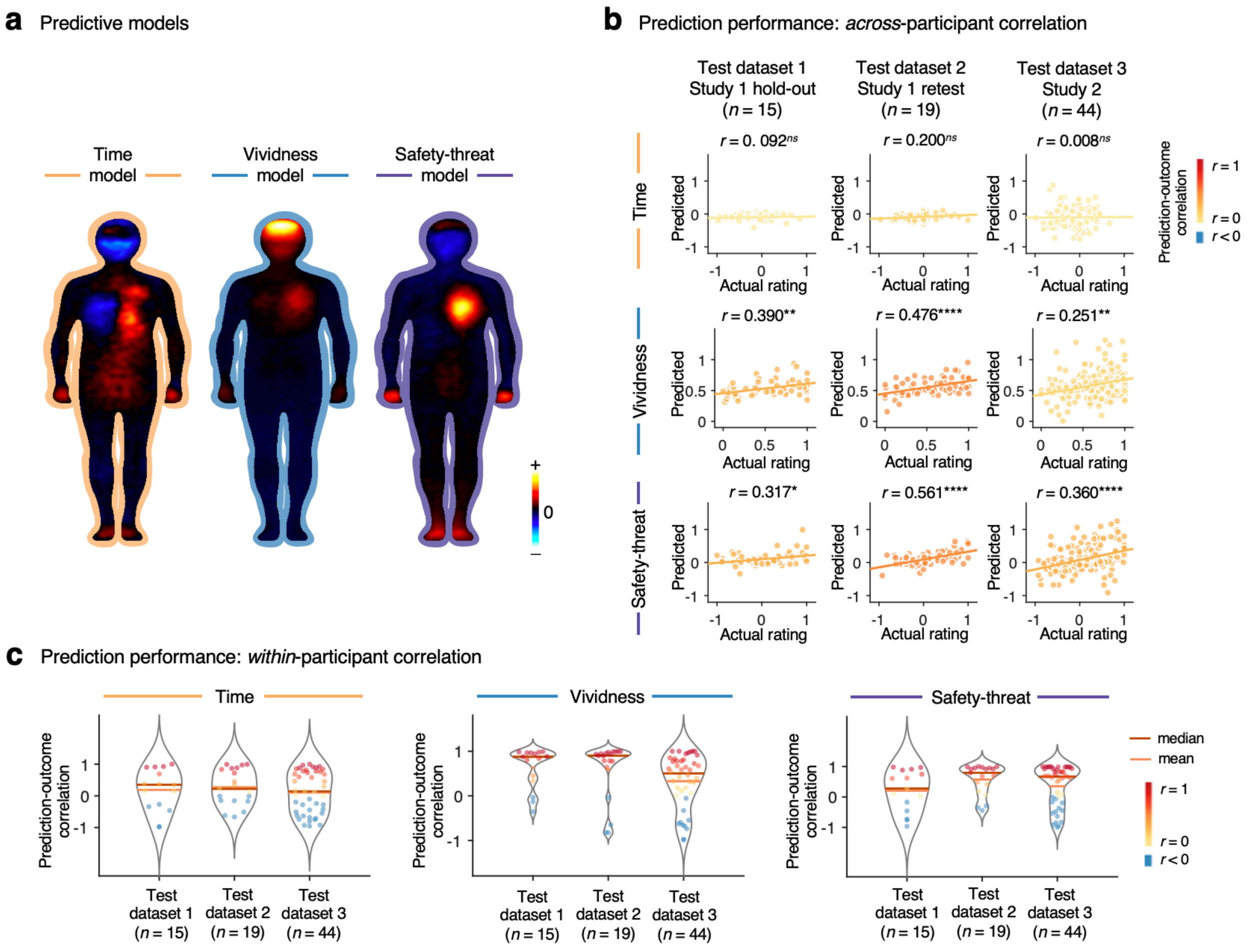


**Supplementary Fig. 5 (related to Fig. 2). Predictive models of the time, vividness, and safety-threat dimensions. a**, Body map-based predictive models of the time, vividness, and safety-threat dimensions. The warm colors indicate positive predictive weights, and the cool colors indicate negative ones. **b**, Prediction performance based on across-participant correlation. The rows indicate the prediction performance of the time (top), vividness (middle), and safety-threat (bottom) models, and the columns represent different test datasets. From left to right, the plots show the results from 1) the hold-out test dataset of Study 1 ($n$ = 15), 2) the test dataset 2 of Study 1 retest data ($n$ = 19, the overlapping participants were cross-validated), and 3) the test dataset 3 of Study 2 ($n$ = 44). Scatter plots show the correlation between the actual versus predicted quartile average scores. Each dot represents each quartile's average score of a participant. The line represents the least-square regression line. The colors of dots and lines indicate the level of prediction-outcome correlation. The prediction-outcome correlation was for time, $r$s = 0.008 to 0.200, $p$s = 0.0916 to 0.9165; for vividness, $r$s = 0.251 to 0.476, $p$s = $1.83\times10^{-5}$ to 0.0023; for safety-threat, $r$s = 0.317 to 0.561, $p$s = $2.95\times10^{-7}$ to 0.0145; two-tailed, one-sample t-test, $^{****}p < 0.0001$, $^{**}p < 0.01$, $^{*}p < 0.05$, $^{ns}p > 0.05$. **c**, Prediction performance based on within-participant correlation. Each dot represents the correlation between the actual versus predicted quartile averages for each participant. The dot colors indicate the level of prediction-outcome correlation. The red and orange lines indicate the median and mean of the participant-level prediction performance, respectively. The averaged correlations between the predicted and actual content dimension scores ranged from 0.106 to 0.270 for the time model, 0.327 to 0.633 for the vividness model, and 0.202 to 0.570 for the safety-threat model, bootstrap tests with 10,000 iterations (please see **Supplementary Table 2** for the details).

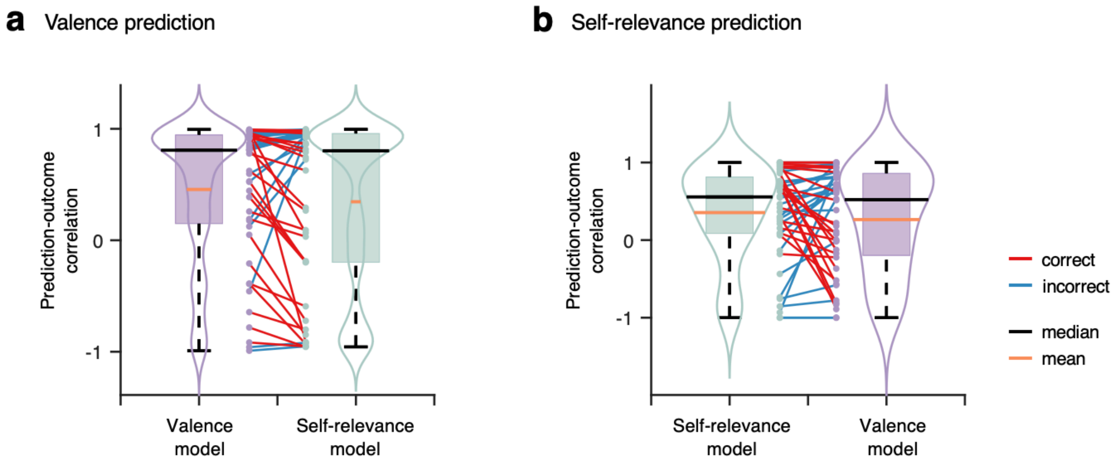


**Supplementary Fig. 6 (related to Fig. 2). Cross-prediction performance of the valence and self-relevance models.** To test the specificity of the models, we conducted the cross-prediction using the test dataset 3 of Study 2 ($n$ = 44). **a,** The plot shows the prediction performance of the valence and self-relevance models predicting the quartile averages of actual valence scores. We show the distributions of the within-participant correlation coefficient using violin and box plots. The boundary of the boxplot was determined by the first and the third quartiles, and the whiskers are marked at the highest and lowest values. Each dot represents the correlation between the actual versus predicted quartile averages for each participant. The red lines indicate the higher prediction performance of the valence model than the self-relevance model for the valence prediction (correct), and the blue lines indicate the lower prediction performance of the valence model than the self-relevance model (incorrect). **b**, The prediction results of the self-relevance using the self-relevance and valence models. The red lines indicate the higher prediction performance of the self-relevance model than the valence model for the self-relevance prediction (correct), and the blue lines indicate the lower prediction performance of the self-relevance model than the valence model (incorrect).

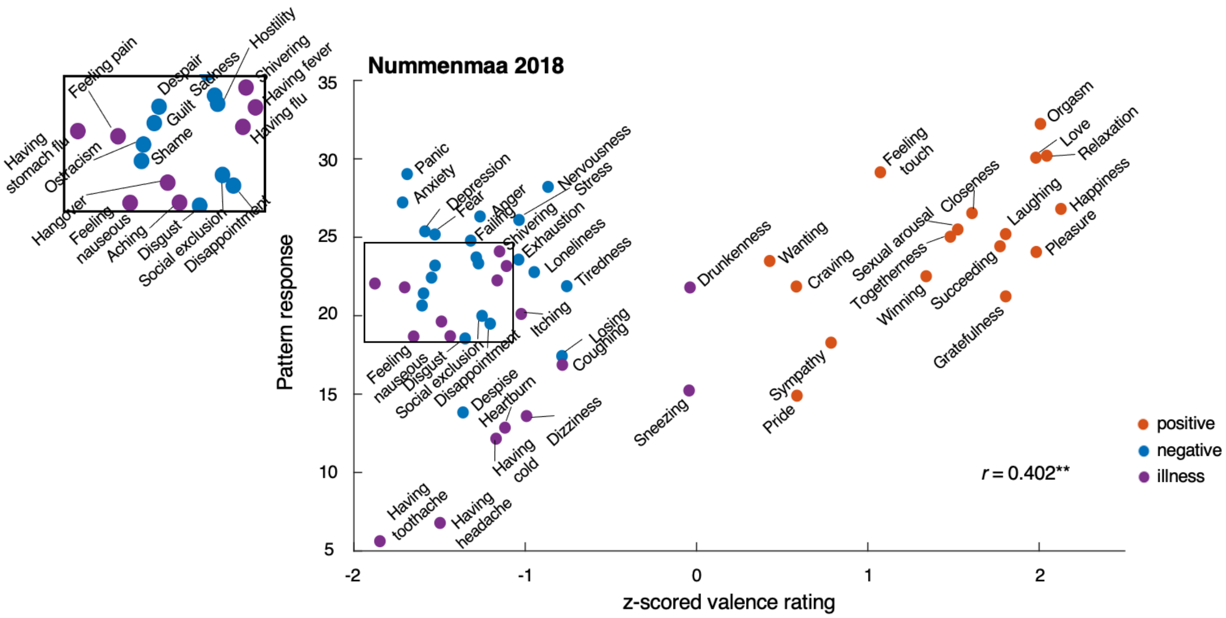


**Supplementary Fig. 7 (related to Fig. 3). Testing results of the valence model on the data from Nummenmaa et al. (2018)[1] with all labels.** The scatter plots show the actual z-scored valence ratings against pattern responses, which were calculated with the dot product of our model and the body maps from the studies. The dot colors indicate the clusters of emotions and feelings according to Nummenmaa et al. (2018)[1]. The inset shows the enlarged version of the data with labels due to the lack of space for the labeling. $^{**}p < 0.01$, two-tailed, one-sample t-test.

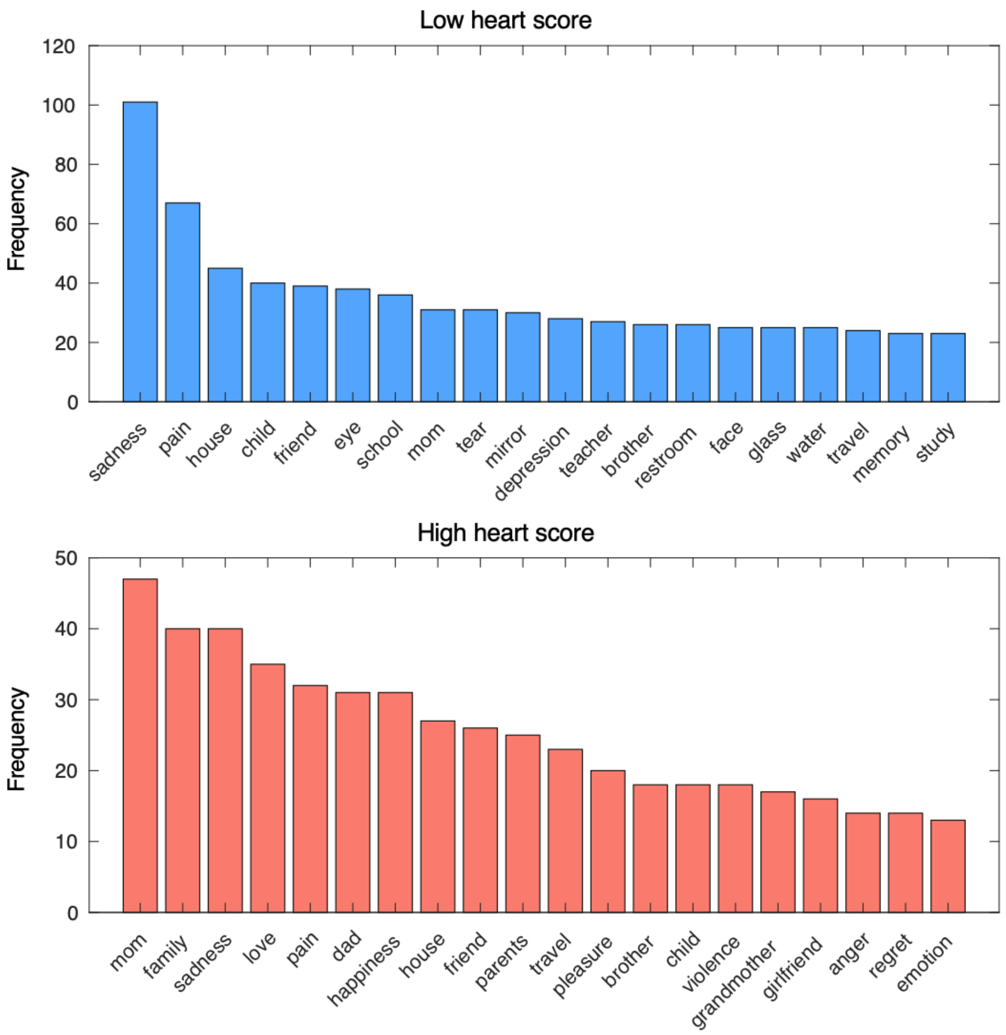


**Supplementary Fig. 8 (related to Fig. 4). Word frequency of high versus low heart cluster score trial groups.** The plots show the top 20 words for each trial group. The data are from Study 1, 50 participants (see **Methods** for the detail of participants).

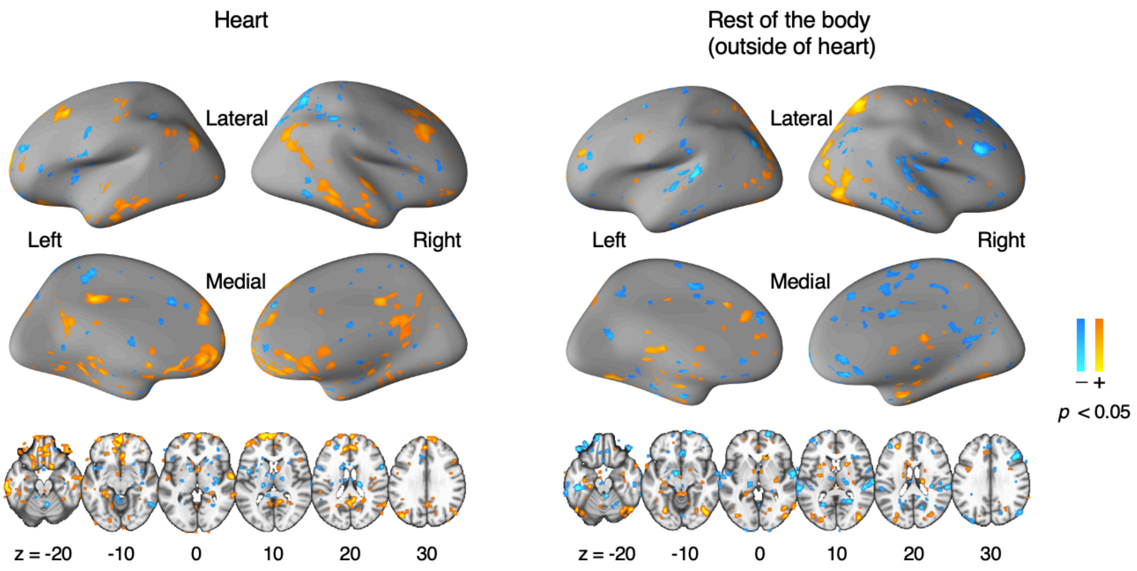


**Supplementary Fig. 9 (related to Fig. 4). Heart cluster-related GLM map with the scores of remaining body parts as a covariate.** The brain maps on the left panel indicate the heart cluster-related GLM maps after controlling for the rest of the body scores (i.e., outside of the heart cluster). The results were largely consistent with the GLM results without the covariate. The brain maps on the right panel show the GLM maps of the covariate (i.e., the remaining body part scores after removing the heart cluster). We thresholded the maps at uncorrected voxel-wise $p < 0.05$.

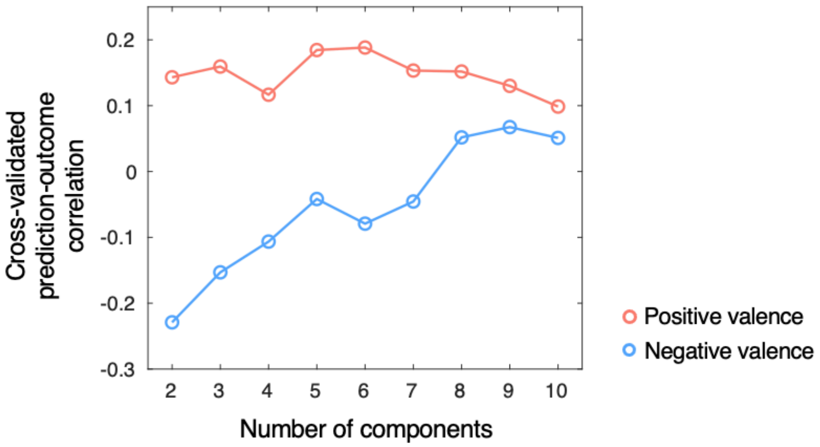


**Supplementary Fig. 10. Body map-based predictive modeling of positive and negative valence.** We trained the positive and negative valence models separately by dividing the training data (participant $n$ = 29) into the trials with positive (≥ 0) versus negative (< 0) valence trials. We used principal component regression with leave-one-subject-out cross-validation. The plot shows the cross-validated prediction-outcome correlation for different numbers of principal components (PCs) and for the two models.

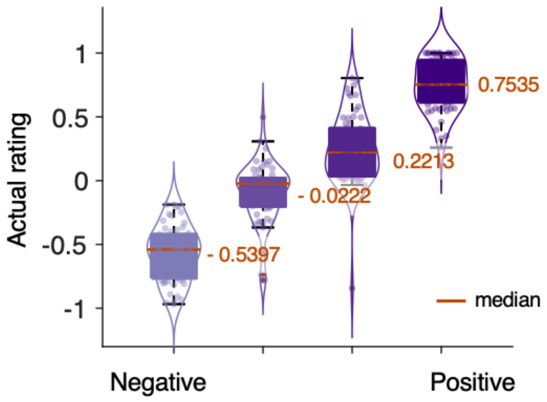


**Supplementary Fig. 11. Distribution of quartile valence scores.** The violin and box plots show the distribution of the quartile valence scores (participant $n$ = 62, Study 1). The boundary of the boxplot was determined by the first and the third quartiles, and the whiskers are marked at the highest and lowest values within the median ± 1.5 interquartile range. Each dot represents the average rating score of a participant for each quartile, and the outliers outside of the range are shown as red dots.

**Supplementary Table 1. Prediction performance: across-participant prediction-outcome correlation**

| Models | Train dataset | | Test dataset 1 | | Test dataset 2 | | Test dataset 3 | |
|---|---|---|---|---|---|---|---|---|
| | *r* | *p* | *r* | *p* | *r* | *p* | *r* | *p* |
| Valence | 0.432 | 0.0000 | 0.422 | 0.0010 | 0.623 | 0.0000 | 0.428 | 0.0000 |
| Self-relevance | 0.359 | 0.0001 | 0.423 | 0.0008 | 0.513 | 0.0000 | 0.297 | 0.0001 |
| Time | 0.026 | 0.7856 | 0.092 | 0.4954 | 0.200 | 0.0916 | 0.008 | 0.9165 |
| Vividness | 0.445 | 0.0000 | 0.390 | 0.0023 | 0.476 | 0.0000 | 0.251 | 0.0011 |
| Safety-threat | 0.202 | 0.0349 | 0.317 | 0.0145 | 0.561 | 0.0000 | 0.360 | 0.0000 |

*Note.* The *p*-values were obtained from two-tailed one-sample *t*-tests for Pearson's correlation.

**Supplementary Table 2. Prediction performance: within-participant prediction-outcome correlation**

| Models | Train dataset | | Test dataset 1 | | Test dataset 2 | | Test dataset 3 | |
|---|---|---|---|---|---|---|---|---|
| | *r* | *p* | *r* | *p* | *r* | *p* | *r* | *p* |
| Valence | 0.541 | 0.0000 | 0.308 | 0.0641 | 0.662 | 0.0000 | 0.456 | 0.0000 |
| Self-relevance | 0.586 | 0.0000 | 0.679 | 0.0000 | 0.651 | 0.0000 | 0.355 | 0.0001 |
| Time | 0.271 | 0.0212 | 0.189 | 0.2556 | 0.270 | 0.0322 | 0.106 | 0.2722 |
| Vividness | 0.643 | 0.0000 | 0.633 | 0.0000 | 0.588 | 0.0000 | 0.327 | 0.0003 |
| Safety-threat | 0.392 | 0.0020 | 0.202 | 0.2353 | 0.570 | 0.0000 | 0.350 | 0.0005 |

*Note.* The *p*-values were from bootstrap tests with 10,000 iterations.
